# 3D bioengineered neural tissue generated from patient-derived iPSCs develops time-dependent phenotypes and transcriptional features of Alzheimer’s disease

**DOI:** 10.1101/2022.07.21.501004

**Authors:** Selene Lomoio, Ravi S. Pandey, Nicolas Rouleau, Beatrice Menicacci, WonHee Kim, William L. Cantley, Philip G. Haydon, David A. Bennett, Tracy L. Young-Pearse, Gregory W. Carter, David L. Kaplan, Giuseppina Tesco

## Abstract

**Background:** Current models to study Alzheimer’s disease (AD) include cell cultures and animal models. Human diseases, however, are often poorly reproduced in animal models. Developing techniques to differentiate human brain cells from induced pluripotent stem cells (iPSCs) provides a novel approach to studying AD. Three-dimensional (3D) cultures to model AD are represented by organoids, neurospheroids, and scaffold-based cultures. Some AD-related phenotypes have been identified across 3D models [1]. However, to our knowledge, none of these studies could recapitulate several AD-related hallmarks in one single model and establish a temporal relation among them. Furthermore, to date, the transcriptomic features of these 3D models have not been compared with those of human AD brains. These data are, in our opinion, key to understanding the pertinency of these models for studying AD-related pathomechanisms over time.

**Methods:** We developed a 3D bioengineered model of iPSC-derived neural tissue that combines a porous scaffold composed of silk fibroin protein with an intercalated collagen hydrogel to support the growth of neurons and glial cells into complex and functional networks. This biomaterial scaffold, designed to match the mechanical properties of brain tissue, can support 3D neural cultures for an extended time without necrosis, a fundamental requisite for aging studies.

We have optimized our protocol by seeding neural precursor cells (NPCs) into these scaffolds. NPC-derived cultures were generated from iPSC lines obtained from two subjects carrying the familial AD (FAD) APP London mutation, two well-studied control lines, and an isogenic control. Cultures were analyzed at 2 and 4.5 months.

**Results:** An elevated Aβ42/40 ratio was detected in conditioned media from FAD cultures at both time points, as previously reported in 2D cultures derived from the same FAD lines. However, extracellular Aβ42 deposition and enhanced neuronal excitability were observed in FAD culture only at 4.5 months. The increased excitability of FAD cultures correlated with extracellular Aβ42 deposition but not with soluble Aβ42/40 ratio levels, as they were similar at both time points. These data suggest that extracellular Aβ deposition may trigger enhanced network activity. Notably, neuronal hyperexcitability has been described in AD patients early in the disease. Transcriptomic analysis revealed the deregulation of multiple gene sets in FAD samples. Notably, such alterations were similar to those observed in human AD brains in a large study that performed a co-expression meta-analysis of harmonized data from Accelerating Medicines Partnership for Alzheimer’s Disease (AMP-AD) across three independent cohorts.

**Conclusions:** Our 3D tissue model supports the differentiation of healthy iPSC-derived cultures in a porous silk-collagen composite sponge with an optically clear central region. This design facilitates nutrient delivery to meet the metabolic demand of long-term cultures. These data provide evidence that our bioengineered model from patient-derived FAD iPSCs develops time-dependent AD-related phenotypes and establishes a temporal relation among them. Furthermore, FAD iPSC-derived neuronal tissue recapitulates transcriptomic features of AD patients. Thus, our bioengineered neural tissue represents a unique tool to model AD-related pathomechanisms over time, with several advantages compared to the existing models.

## Background

Alzheimer’s disease (AD) is a progressive neurodegenerative disorder characterized by memory impairments and cognitive deterioration. It is one of the most common forms of neurodegeneration and accounts for approximately 70% of all cases of dementia. The key pathological hallmarks of AD are extracellular senile plaques composed of neurotoxic amyloid-beta (Aβ) peptide and intracellular neurofibrillary tangles (NFT) composed of hyperphosphorylated tau [2]. Clinical studies show that the most critical risk factors for AD are aging and family history of the disease. Approximately 5% of the population over 65 is affected by AD, and its prevalence doubles with every five years of increasing age [3]. Less than 1% of AD cases occur in families where the disease is inherited in a fully penetrant, autosomal dominant manner, with early (< 65) onset. Three early-onset familial AD (FAD) genes have been identified to date: the amyloid precursor protein (APP) gene, the presenilin 1 (PSEN1) gene, and the presenilin 2 (PSEN2) gene. Together, these genes can harbor any of > 400 mutations (https://www.alzforum.org/mutations) that account for roughly 5-10% of autosomal dominant FAD.

Current models to study AD include cell cultures and animal models. Human diseases, however, are often poorly reproduced in animal models [4]. A significant limitation of using rodents to model AD is the increasing evidence for marked differences between humans and rodents in glial gene expression, in contrast to the relatively conserved neuronal gene expression across mammals [5-7]. The development in the past decade of techniques to generate human-induced pluripotent stem cells (iPSCs) and differentiate them into different brain cell types, together with advances in genome editing, has provided a novel model to study AD and other neurodegenerative disorders [1, 8]. Neural cultures derived from patients or isogenic mutant iPSCs recapitulate at least some of the phenotypes observed in the AD brain [1, 8, 9]. However, conventional two-dimensional (2D) culture systems fail to recapitulate the diversity and maturation of multiple cell types and their interaction under physiological and pathological conditions. For example, removing secreted Aβ with media changes most likely prevents the deposition of amyloid aggregates in 2D cultures [1]. To date, three-dimensional (3D) cultures to model AD are represented by organoids, neurospheroids, and scaffold-based 3D cultures (for review, see [1]). Some AD-related phenotypes have been identified across 3D models [1]. However, to our knowledge, none of these studies could recapitulate several AD-related hallmarks in one single model and establish a temporal relation among them. Furthermore, to date, no studies have compared the transcriptomic features of human 3D models to those of human AD brains. These data are, in our opinion, key to understanding the pertinency of these models for studying AD-related pathomechanisms over time.

We developed a bioengineered model of iPSC-derived neural tissue [10]. This biomaterial scaffold design was adapted from our previous tissue model generated using primary embryonic rodent brain cells [11] and human-induced neural stem cells [12, 13] to match the mechanical properties of native brain tissue. Our silk-collagen-based “donut” scaffolds can support compartmentalized, 3D neural-like tissues for over a year without necrosis [14]. More importantly, our cultures develop networks and action potentials (APs) typical of fully mature neurons [10]. The architecture of the scaffolds was optimized to meet the metabolic demand of high-density cultures in terms of free diffusion of nutrients and oxygen, a fundamental requisite for long-term cultures and aging-related studies.

To support that our bioengineered tissue model can reproduce AD-related phenotypes, we have generated cultures from iPSC lines obtained from two subjects carrying the FAD mutation APP V717I (also known as London), hFAD-1, and hFAD-2 (Clone A and B) [15]. As controls, we used two well-studied lines, YZ1 and BR24 [9, 15]. Furthermore, we generated the isogenic control of the hFAD-2 Clone B line by correcting the APP V717I mutation to wild type (C-hFAD-2 Clone B). FAD and control cultures were analyzed at two time points (2 and 4.5 months). An elevated Aβ42/40 ratio was detected in FAD cultures at both time points, as previously reported in 2D cultures derived from the same FAD lines [15].

Additionally, extracellular Aβ42 deposition and enhanced neuronal excitability were observed in FAD culture at 4.5 months. Remarkably, neuronal hyperexcitability has been described in both FAD and late-onset AD (LOAD) patients at the early stage of the disease [16]. Gene expression analysis was performed to identify the specific target genes which exhibit significantly increased or decreased expression in FAD lines compared to controls at both time points. Multiple gene sets were deregulated in FAD samples at 2 months. The difference was maintained up to 4.5 months. Several of the pathways identified are implicated in neurotransmitter release/reuptake and synthesis/storage, neural connectivity, structural stability of the cytoskeleton, axonal and dendrite structure, and vesicle trafficking regulation. Such alterations in gene expression were similar to those observed in human AD brains in a large study that performed a co-expression meta-analysis of harmonized Accelerating Medicines Partnership for Alzheimer’s Disease (AMP-AD) data across three independent cohorts [17]. These data provide compelling evidence that our 3D cultures reproduce some time-dependent phenotypes and transcriptomic features observed in AD patients. Thus, our bioengineered neural tissue represents a valuable model for studying AD-related pathomechanisms over time.

## Results

### A 3D bioengineered silk-collagen neural model

We have previously designed and validated a 3D bioengineered neural model that combines a porous scaffold composed of silk fibroin with an intercalated hydrogel to allow neurons and glial cells to grow into complex and functional networks [11, 13, 18]. Each element of this model was designed to be tunable, from the pore sizes in the scaffold biomaterial [19] to the hydrogel composition, mechanical properties, and incorporated cell types [14]. We demonstrated that iPSCs incorporated into the scaffolds could directly differentiate into neurons and glial cells and remain functionally active for several months [10].

We have optimized our protocol for this study by seeding neural precursor cells (NPCs) instead of iPSCs. This improved protocol offers several advantages: NPCs can be expanded, frozen, banked, and subsequently differentiated. Having NPC stocks derived from multiple subjects allows synchronization of their differentiation and minimizes experimental variability. We have successfully generated NPCs from various iPSC lines using an embryoid body (EB)-based protocol [15, 20-22] that seeks to recapitulate embryonic differentiation. In the absence of exogenous patterning factors, the default path of this protocol is to generate forebrain neurons of cortical fates [23].

For the success of this experimental strategy, quality control parameters are associated with iPSC cultures [24]. iPSCs are susceptible to cell culture stresses and prone to genomic instability. Morphology is an obvious indicator of the iPSC health [24]. In our practice, karyotyping, potency, and mycoplasma testing have been routinely performed in all iPSC lines used for this study (Fig. 1A-B, Fig. S1).

**Figure 1.**
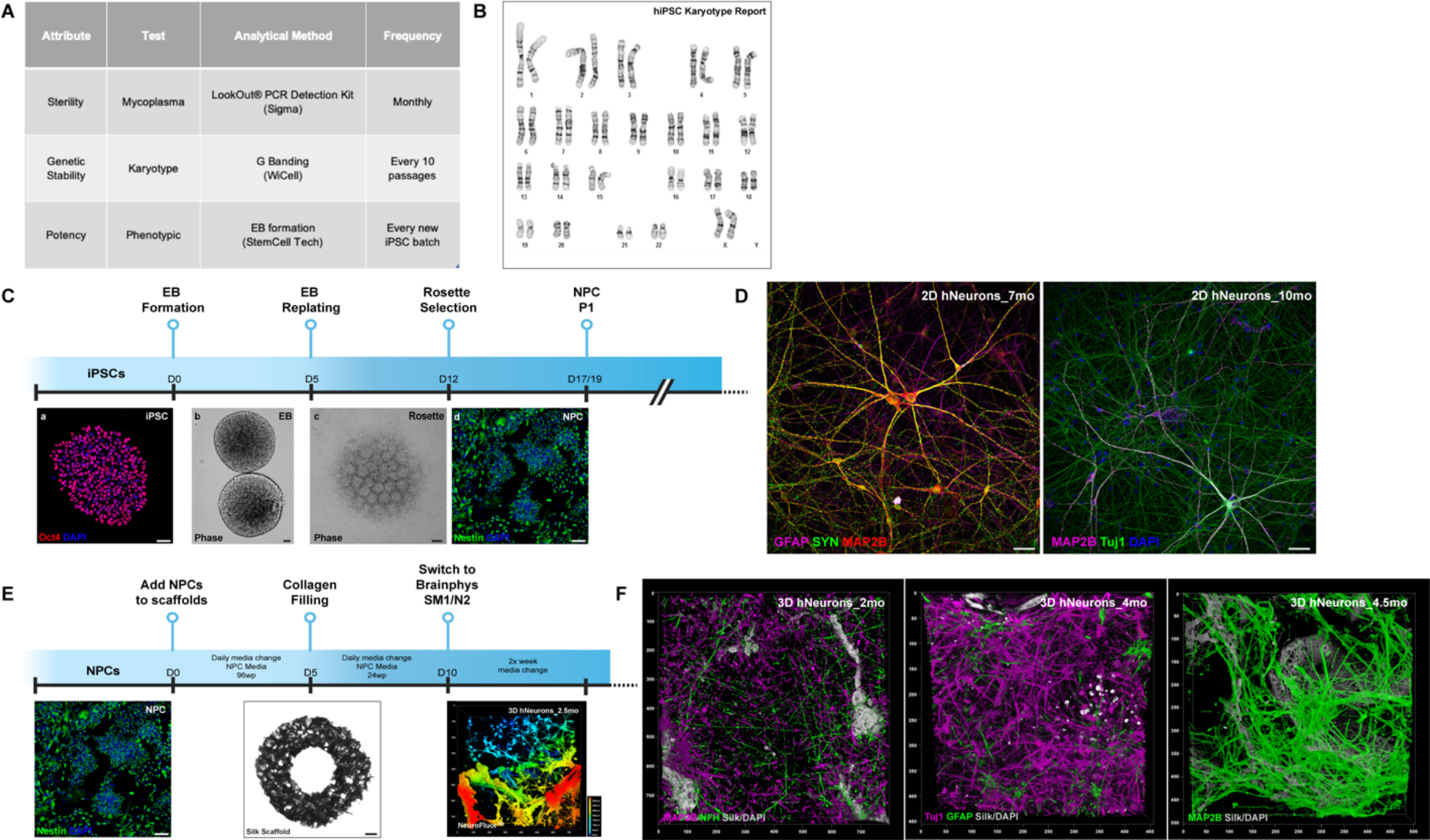
A 3D bioengineered silk-collagen neural model. (A) Quality tests performed on iPSCs used in this study. (B) Representative karyotype report. Reports for all the lines are in Fig. S1. (C) Schematic representation of the differentiation timeline to generate NPCs. (a) Representative iPSC colony stained for Oct4 (red) and DAPI (blue). Scale bar: 40um. Confocal z-stack. Leica SP8 HC PL APO CS2 20x/0.75 IMM. 1.5x Zoom. EB (b) and rosettes (c) phase-contrast images. Scale bar: 20um. Nikon TE300 CFI PL FL DL 4x/0.13 and 10x/0.30. (d) NPC cultures stained for nestin (green) and DAPI (blue). Scale bar: 40um. Confocal z-stack. Leica SP8 HC PL APO CS2 20x/0.75 IMM. 1x Zoom. (D) 2D 7-months-old hNeurons stained for GFAP (magenta), SYN (green), and MAP2B (red), and 10-months-old hNeurons stained for MAP2B (magenta), Tuj1 (green), and DAPI (blue). Scale bar: 50um. Confocal z-stack. Leica SP8 HC PL APO CS2 20x/0.75 IMM. 0.75x Zoom. (E) Schematic representation of the timeline to differentiate hNeurons in silk-collagen scaffolds from NPCs. Silk scaffold phase-contrast image. Scale bar: 500um. Z-stack. Stitched image. Zeiss CD7 PL APO 5x/0.35. Magnification changer 0.5x. 3D reconstruction of hNeurons grown in a scaffold for 2.5 months and stained with NeuroFluor™ NeuO. Leica SP8 HC PL APO CS2 20x/0.75 IMM. 0.75x Zoom. (F) 3D reconstruction of hNeurons grown in a scaffold for 2 months (stained for MAP2B (magenta), NFH (green)), 4 months (stained for Tuj1 (magenta), GFAP (green)) and 4.5 months (stained for MAP2B (green)). For all the images DAPI and silk autofluorescent signals are in grey. Leica SP8 HC PL APO CS2 20x/0.75 IMM. 0.75x Zoom.

Figure 1C illustrates the timeline schematic for the protocol, in which well-defined and compact iPSC colonies expressing pluripotency markers like the octamer-binding transcription factor 4 (Oct4) are differentiated to NPCs over approximately 20 days. The milestones of this approach are the formation of EBs, three-dimensional cell aggregates that mimic some structures of the developing embryo. They can differentiate into all three germ layers [25], followed by neural rosette maturation that can be identified morphologically by their characteristic appearance as round clusters of primitive neuroepithelial cells with apicobasal polarity [26] (Fig. 1C).

According to previous work, nestin-positive NPCs can be robustly passaged, frozen, thawed, and differentiated into functional neurons. NPCs could be typically passaged more than ten times in vitro without accumulating karyotype abnormalities [26]; however, in agreement with other studies [26, 27], we found that beyond ten passages, NPCs show increased cellular heterogeneity and lower differentiation efficiency. Thus, experiments were conducted on mycoplasma-free passage-matched NPC populations between passages two and six.

When differentiated in 2D on a poly-ornithine/laminin substrate using BrainPhys media supplemented with SM1 and N2, NPCs can efficiently differentiate into human neurons (hNeurons) expressing characteristic markers of a mature neuronal population such as the microtubule-associated protein 2B (MAP2B), the neuron-specific class III beta-tubulin (Tuj1), and synaptophysin (SYN) for extended periods. Glial fibrillary acidic protein (GFAP)-positive cells are also observed, indicating the presence of astrocytic-like cells (Fig. 1D). NPCs have already been described as having dual lineage potential and can differentiate into 70-80% neural populations of Tuj1-positive neurons and 20-30% GFAP-positive astrocytes [26].

For this study, NPCs have been seeded into silk scaffolds and cultured according to the timeline in Fig. 1E. Our data indicate that NPCs integrate into the silk-collagen scaffold model, expand (NeuroFluor NeuO live staining, Fig. 1E), and differentiate into healthy neurons and astrocytic-like cells, as indicated by IF imaging (Fig. 1F). Cultures express neuronal markers such as neurofilament heavy chain (NFH), MAP2B, and Tuj1 together with GFAP (Fig. 1F). They develop an intricated three-dimensional network extending deep into the silk pores and the supporting collagen, suggesting high connectivity among growing cells (Fig. 1E, F). Cultures generated with this approach supported by BrainPhys media supplemented with SM1 and N2 can survive in culture for extended periods.

### Aβ42/40 ratio and extracellular Aβ42 deposition are elevated in FAD cultures

To explore whether our bioengineered cultures could be valuable for modeling AD, we generated cultures from extensively characterized iPSC lines [15] from a father (hFAD-1) and a daughter (hFAD-2), each harboring the FAD mutation APP V717I [15] (Fig. 2A). The father was 57 years old and diagnosed with AD before the biopsy. In contrast, the daughter was asymptomatic at age 33. The two clones from the daughter (hFAD-2 Clone A and Clone B) displayed a normal euploid karyotype (Fig. S1), while the clone from the father (hFAD-1) revealed a standard chromosome number but a balanced (t(1;12)(q42.3;q21.2)) translocation in all cells analyzed (Fig. S1). The fibroblasts obtained from this subject displayed the same abnormal karyotype, suggesting a preexisting abnormality that did not arise during the reprogramming [15]. This karyotypic alteration didn’t compromise the differentiation potential for the line.

**Figure 2.**
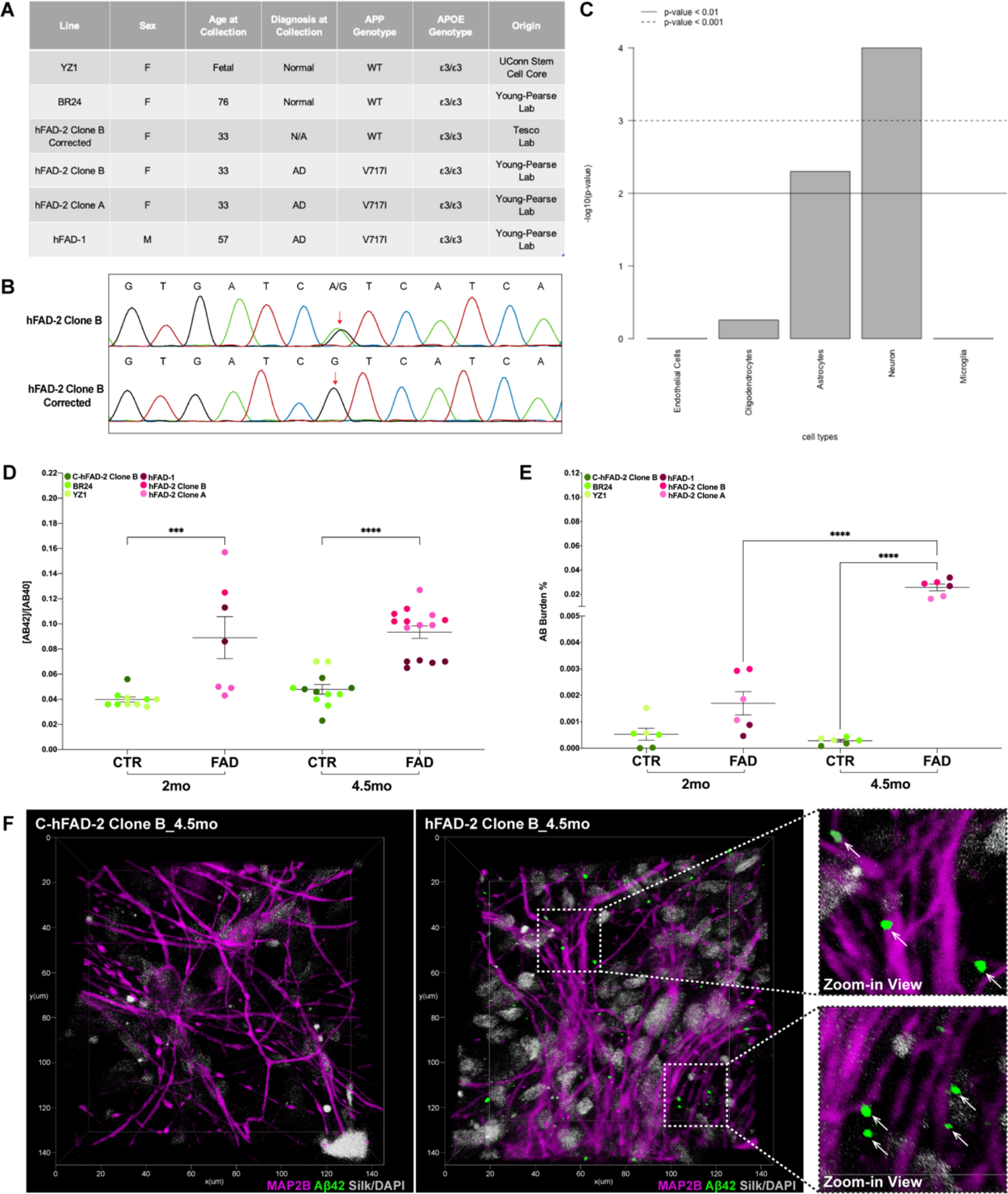
Aβ42/40 ratio and extracellular Aβ42 deposition are elevated in FAD cultures. (A) Table reporting iPS lines used across the study. (B) Sequencing traces of hFAD-2 Clone B and its isogenic control after reversion of the APP V717I mutation (C-hFAD-2 Clone B). (C) nSolver Cell Enrichment analysis bar plot. (D) Aβ42/40 MSD analysis quantification in conditioned media. FAD cultures (hFAD-1, hFAD-2 Clone A, hFAD-2 Clone B) and controls (C-hFAD-2 Clone B, BR24, YZ1) plotted as groups are provided at 2 and 4.5 months. CTR 2mo: n=10, FAD 2mo: n=7, CTR 4.5mo: n=12, FAD 4.5mo: n=15. One-way ANOVA, Tukey’s post-hoc test. (E) Amyloid burden quantification (%). FAD cultures (hFAD-1, hFAD-2 Clone A, hFAD-2 Clone B) and controls (C-hFAD-2 Clone B, BR24, YZ1) plotted as groups are provided at 2 and 4.5 months. n=6. One-way ANOVA, Tukey’s post-hoc test. (E) Representative 3D reconstructions of hNeuron culture (4.5 months) derived from C-hFAD-2 Clone B and hFAD-2 Clone B stained for MAP2B (magenta) and Aβ42 (green). DAPI and silk autofluorescent signals are in grey. Zoom-in view: green Aβ42 extracellular depositions (white arrows) interspersed among hNeurons (MAP2B, magenta). Leica Falcon SP8 HC PL APO CS2 20x/0.75 IMM. 3.5x Zoom.

The following lines were selected as controls (Fig. 2A). YZ1 [15], derived from fetal lung fibroblasts, BR24 (Religious Order Study (ROS) or Rush Memory and Aging Project (MAP) [28]), a cognitively unimpaired woman at the time of biopsy, 76 years of age [9]. Participants in the ROS and MAP projects range from 50 to over 100 years and are free of dementia at the enrollment [9, 28]. Follow-up assessments indicated that the BR24 subject died at the age of 90, showing no signs of cognitive decline or AD pathology. We also generated an isogenic control for the hFAD-2 Clone B line (Fig. 2B) using genome-editing technology by reverting the APP V717I mutation to wild type (C-hFAD-2 Clone B). Potential off-target cuts were identified using the Cas-OFF finder (http://www.rgenome.net/cas-offinder). The top 8 sites were amplified, and Sanger sequenced to verify the status of these potential off-targets. None of the predicted sites showed evidence of off-targeting cutting. No karyotype clonal abnormalities were identified among the control lines (Fig. S1).

To better understand the nature of our cultures and the reproducibility of the differentiation approach from different iPSC lines and clones, we next turned to NanoString Technologies^®^ for measuring gene expression at 2 and 4.5 months. Data were analyzed by ROSALIND^®^ (https://rosalind.onramp.bio), with a HyperScale architecture developed by ROSALIND^®^, Inc. (San Diego, CA, USA), and the nCounter^®^ Advanced Analysis protocol. Housekeeping probes (9) used for normalization were selected based on the geNorm algorithm implemented in the NormqPCR R library.

We used the nCounter^®^ Cell Type Profiling Module (as described in [29]) to assess the abundance of various cell populations (Fig. 2C). The method quantifies cell populations using marker genes expressed stably and precisely in given cell types. While markers for endothelial cells, oligodendrocytes, astrocytes, neurons, and microglia have been screened, only scores for neurons and astrocytes reached a statistically significant threshold (p-value <0.001 and <0.01, respectively, Fig. 2C), indicating that these two cell types are the main constituents of our cultures. Exploiting NanoString normalized expression analysis, we compared the expression of specific groups of commonly expressed genes in iPSC-derived neural cultures (Fig. S2) based on previous studies [15, 20, 23, 30-32] between different lines, genotypes, time points, and biological replicates. In agreement with previous observations [15, 20], normalized expression analysis identified the enrichment of a vast array of glial (GFAP, ALDH1L1, S100B, etc.), pan-neuronal and synaptic markers (HOMER1, TBR1, STX1A, etc.) with most neurons exhibiting an excitatory profile, together with some progenitor markers (NES, CCND1) being expressed up to 4.5 months post-seeding. As expected, based on the differentiation method implemented, genomic analysis suggests that our culture acquired a cortical layer fate as indicated by the expression of genes (TBR1, DLX1, DLX2, etc.) involved in mammalian neocortical development and/or expressed in a layer-specific manner in the human neocortex (Fig. S2) [33]. Also, AD-related genes were expressed in our 3D cultures across all analyzed conditions. Differential expression analysis did not reveal any significant variance between lines and genotypes for this set of genes, suggesting that the cultures are homogeneously patterned, and that the differentiation approach is highly reproducible.

To assess whether our cultures can recapitulate some AD-related phenotypes, soluble Aβ quantification (MSD: Meso Scale Discovery Assay, Aβ Peptide Panel 1-4G8) was performed on 24-hour-conditioned media (Fig. 2D). In agreement with previous reports [15, 23, 34, 35], the Aβ42 over Aβ40 ratio was found to be increased in FAD cultures compared to control ones at both time points (CTR 2mo: 0.040±0.002 vs. FAD 2mo: 0.089±0.017, p=0.0003; CTR 4.5mo: 0.048±0.004 vs. FAD 4.5mo: 0.093±0.005, p<0.0001). The Aβ42/40 ratio calculated for the isogenic C-hFAD-2 Clone B control was comparable to that of other control lines, supporting the idea that the observed increase in the hFAD-2 Clone B depended on the APP V717I mutation.

Next, we performed IF using an antibody specific for Aβ42 (validated in 5XFAD:BACE1^-/-^ brain sections [36]) and calculated the amyloid burden for CTR and FAD cultures at both time points (Fig. 2E). Confocal imaging and 3D reconstruction revealed the presence of Aβ42-positive extracellular aggregates interspersed among FAD hNeurons (identified by MAP2B staining, Fig. 2F). Volumetric analysis of the amyloid burden (Fig. 2E) revealed a mild trend for increase starting at 2 months in FAD cultures compared to controls (CTR: 0.00053±0.00022 vs. FAD: 0.00170±0.00044, p=0.9368) with a striking difference among conditions at 4.5 months (CTR: 0.00028±0.00005 vs. FAD: 0.02572±0.00281, p<0.0001). No difference was identified between control groups at the two time points analyzed.

### FAD cultures manifest enhanced neuronal excitability

To determine the electrophysiological properties of the cultures, local field potentials (LFPs) were recorded from single electrodes inserted into the superficial layers of the hydrogel next to silk fibers, adapting a protocol by Du and collaborators [37], where both spontaneous and evoked potentials (EPs) were collected. LFPs, indicative of cellular electrical activations [14], were quantified and compared across all conditions. The spontaneous and evoked activity was observed within all the scaffolds analyzed starting at 2 months, suggesting that our 3D model is populated by functional neurons capable of network activity.

Evidence indicates that LOAD and FAD patients show enhanced excitability, with onset during the early stages of the pathology [38-45]. In our study, FAD cultures displayed comparable levels of spontaneous potentials (Log10 LFP spikes/min) relative to control samples at 2 months (CTR: 0.717±0.130 vs. FAD: 0.872±0.126, p=0.3950); however, by 4.5 months, FAD samples displayed significantly more spontaneous potentials relative to controls (CTR: 0.515±0.106 vs. FAD: 0.832±0.114, p=0.0475). Similarly, FAD samples showed more EPs relative to controls at 4.5 months post-seeding (CTR: 0.736±0.079 vs. FAD: 1.050±0.077, p=0.0064), which was not apparent at 2 months (CTR: 1.040±0.091 vs. FAD: 1.270±0.112, p=0.1069). Our findings agree with previous work that indicates dysregulation of the electrophysiological properties in AD brains accompanied by enhanced network activity. Cognitive impairment in AD patients was assumed to originate from depressed synaptic activity that eventually led to neurodegeneration. However, multiple lines of evidence now indicate that, particularly during AD initial stages, synapse dysfunctions are first induced by neuronal hyperactivity rather than hypoactivity [46-48].

### Differential expression and functional analysis of the NanoString panel in FAD cultures

Transcriptional profiling of NanoString data from our samples was performed by ROSALIND^®^ platform (see method section for details). At 2 months, there were 133 differentially expressed genes (DEGs) (absolute fold change > 1.5 & FDR < 0.05) between FAD and control samples (Fig. 4A). Expression for 38 of those 133 genes was upregulated in FAD samples compared to controls and downregulated for 95 of 133 genes (Fig. 4A). Gene set analysis (GSA) identified increased expression of genes associated with Cytokines, Activated Microglia, and Neuronal Cytoskeleton (directed GSS > 0) (Fig. 4C).

**Figure 3.**
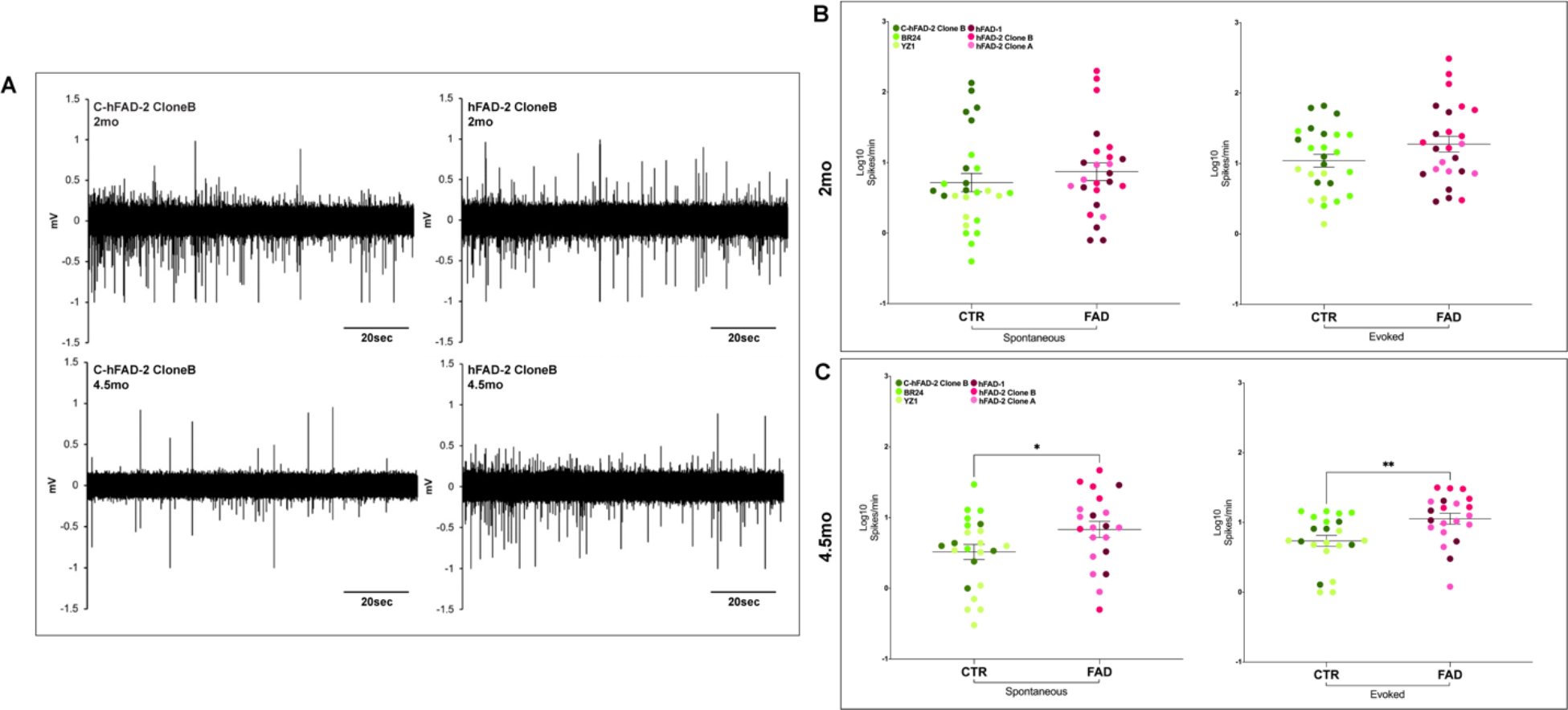
FAD cultures manifest enhanced neuronal excitability. (A) Representative traces of spontaneous LFPs (mV) obtained from 3D cultures derived from a FAD line (hFAD-2 Clone B) and the isogenic control (C-hFAD-2 Clone B) at 2 months and 4.5 months. A 20-sec reference is provided for traces. Quantification of normalized spontaneous and evoked LFP spikes/min (Log10) at 2 months (B) and 4.5 months (C). FAD cultures (hFAD-1, hFAD-2 Clone A, hFAD-2 Clone B) and controls (C-hFAD-2 Clone B, BR24, YZ1) plotted as groups are provided. (B) Spontaneous CTR 2mo n=26, FAD 2mo n=25. Evoked CTR 2mo n=26, FAD 2mo n=25. (C) Spontaneous CTR 4.5mo n=23, FAD 4.5mo n=21. Evoked CTR 4.5mo n=22, FAD 4.5mo n=21. Unpaired t-test.

**Figure 4.**
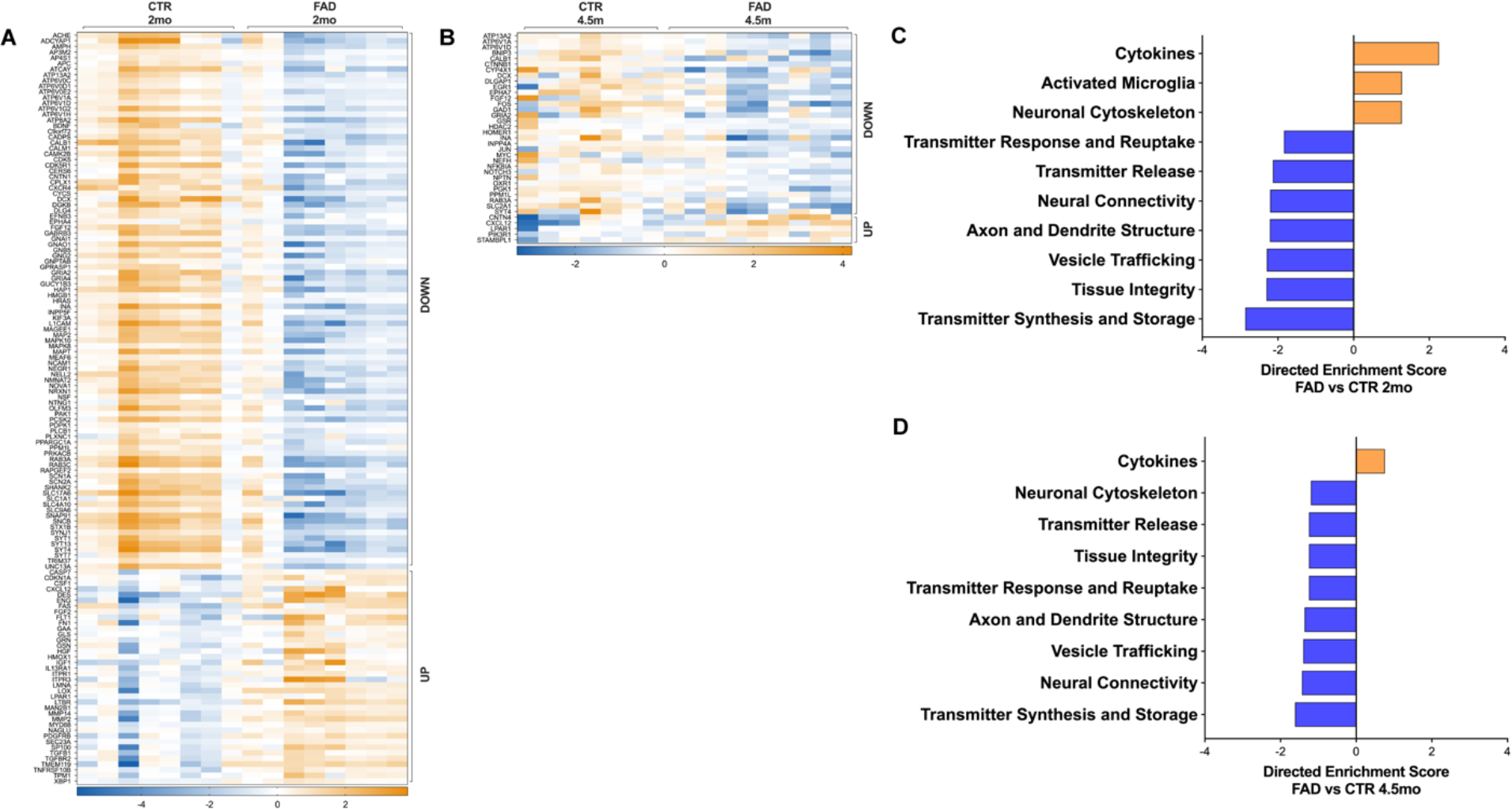
Differential expression and functional analysis of the NanoString panel in FAD cultures. DEGs (absolute fold change > 1.5, FDR < 0.05) in 2-months-old (A) and 4.5-months-old (B) CTR and FAD lines, are shown in the heatmap. GSA of differentially expressed genes in iPSC-derived cultures (FAD vs CTR, ROSALIND^®^ platform, NanoString annotations) at 2 months (C) and 4.5 months (D). Upregulation: directed enrichment score > 0. Downregulation: directed enrichment score < 0. FAD 2mo n=8, CTR 2mo n=8, FAD 4.5mo n=9, CTR 4.5mo n=7.

Based on the differentiation protocol used for this study, we expected our cultures to be populated exclusively by cells derived from the neuroectodermal lineage. So, we were surprised to observe an upregulation of the Activated Microglia pathway in FAD cultures. Follow-up immunofluorescence for TMEM119 (Fig. S3), a well-known microglia marker, followed by confocal microscopy identified the presence of TMEM119 positive cells in the scaffolds interspersed among MAP2 and GFAP-positive cells, indicating the presence of microglia-like cells in our cultures. The cells show a ramified morphology and are intertwined with neuronal processes as described by others (Fig. S3) [49]. These data agree with previous studies that have identified mesoderm-derived progenitors and microglia in cerebral organoids [49, 50].

Further, GSA identified reduced expression of genes associated with multiple other neurodegeneration pathways such as Transmitter Release, Neural Connectivity, and Vesicle Trafficking (directed GSS < 0) (Fig. 4C).

At 4.5 months, there were 38 DEGs (absolute fold change > 1.5 & p < 0.05) between FAD and control samples (Fig. 4B). Expression for 5 of the 38 genes was upregulated in FAD samples compared to controls and downregulated for 33 of 38 genes (Fig. 4B). Gene set analysis identified increased expression of genes associated with Cytokines and reduced expression of genes associated with multiple pathways such as Transmitter Release, Neuronal Cytoskeleton, and Vesicle Trafficking (Fig. 4D).

### FAD cultures reproduce human neurodegeneration signatures

To assess the relevance of our model for AD, we performed a correlation analysis between our NanoString data from FAD cultures and human post-mortem co-expression modules [17, 51]. FAD cultures at 2 months showed significant positive correlations (p<0.05) with AD-associated changes in human co-expression modules in Consensus Cluster B, which include transcripts that are enriched for immune-related pathways in the cerebellum brain region (CBEturquoise) (Fig. 5A). Furthermore, FAD cultures at 2 months showed significant positive correlations (p<0.05) with AD changes in multiple modules in Consensus Cluster C, which include transcripts that are enriched for neuronal-related pathways in the inferior frontal gyrus module (IFG), temporal cortex (TCX), and the parahippocampus gyrus (PHG) brain regions (Fig. 5A). However, these significant positive correlations diminish at 4.5 months (Fig. 5A). Interestingly, FAD cultures at 2 months showed a significant positive correlation (Pearson Correlation = 0.27; p<0.05) with AD ECM organization changes in module TCXblue in Consensus Cluster A but also showed a significant negative correlation (Pearson Correlation = -0.35; p<0.05) with AD changes in the PHGyellow module enriched for similar biological process. However, at 4.5 months of age, FAD cultures exhibited a non-significant but positive correlation with AD changes in the PHGyellow module (Pearson Correlation = 0.11; p > 0.05) and a stronger significant positive correlation with TCXblue module changes in AD (Pearson Correlation = 0.27; p < 0.05) (Fig. 5A). These results suggest that functional changes such as ECM organizations are strengthened with age and represent similar effects to those observed in endpoint AD.

**Figure 5.**
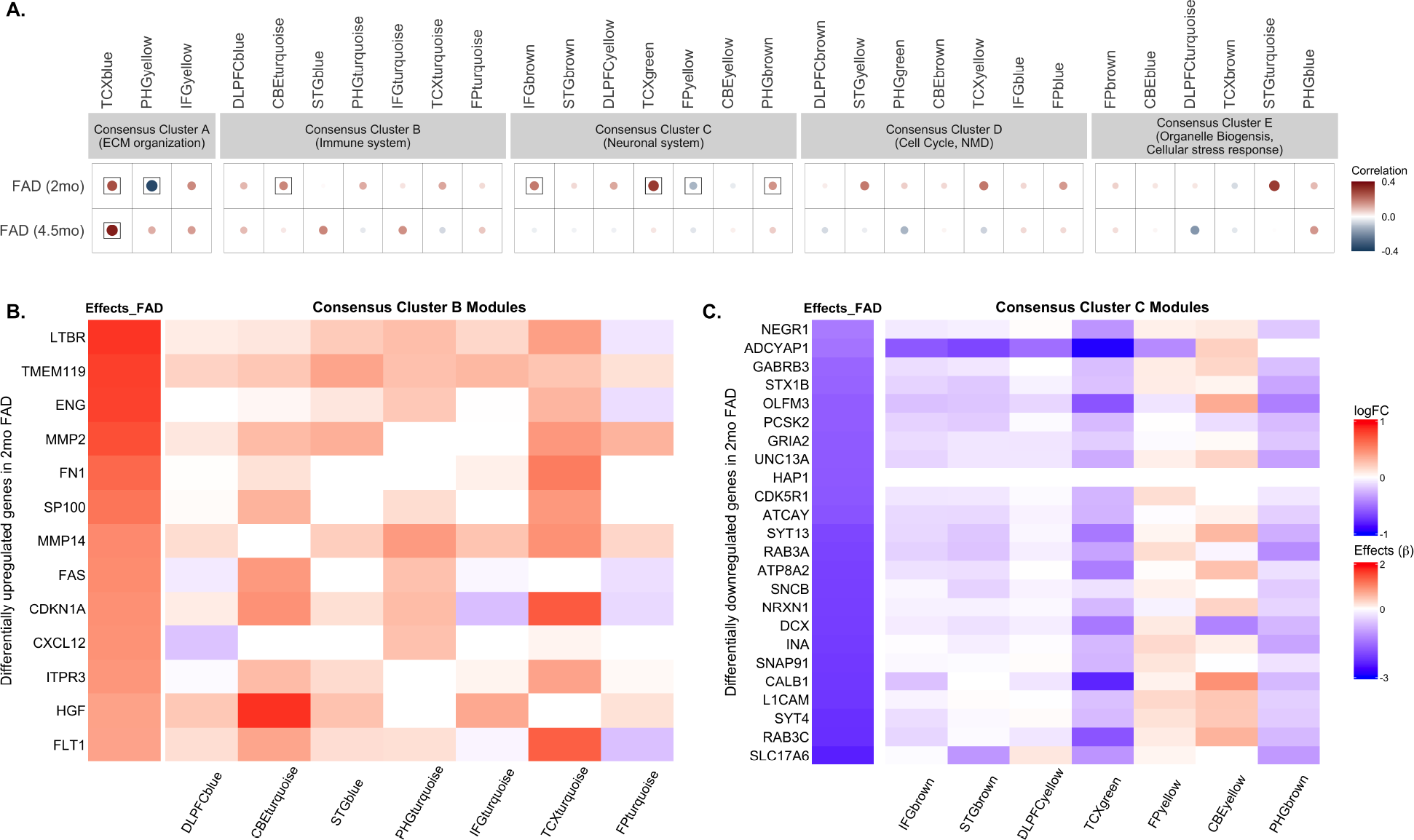
FAD cultures reproduce human neurodegeneration signatures. Correlation analysis between the gene expression profile of FAD cultures and human post-mortem co-expression modules. (A) 2-months-old FAD cultures exhibited a significant positive correlation (p < 0.05) with AD changes in immune-related modules in Consensus Cluster B and neuronal related modules in Consensus Cluster C. Circles within a square correspond to significant (p < 0.05) positive (red) and negative (blue) Pearson correlation coefficients for gene expression changes in FAD lines associated with disease-associated changes in distinct human co-expression modules. Color intensity and size of the circles are proportional to the correlation. (B) Differentially upregulated genes in the 2 months old FAD lines (FDR <0.05, fold change > 1.5), showing increased expression in immune/inflammation associated AMP-AD modules in Consensus Cluster B, are shown in the heatmap. (C) Differentially downregulated genes in the 2 months old FAD lines (FDR <0.05, fold change < 1.5), exhibiting reduced expression in neuronal-associated AMP-AD modules in Consensus Cluster C, are shown in the heatmap. FAD 2mo n=8, CTR 2mo n=8.

Next, we examined the expression profile of differentially regulated genes in the FAD cultures identified by the ROSALIND^®^ platform across human AMP-AD transcriptomic data. We determined that most of the differentially upregulated genes (fold change > 1.5) in 2-month-old FAD cultures showed increased expression in immune/inflammation-associated AMP-AD modules in Consensus Cluster B (Fig. 5B), suggesting increased expression of these genes in multiple human brain regions. Similarly, most of the differentially downregulated genes (fold change < 1.5) in 2 months old FAD cultures showed reduced expression across numerous human brain regions except in the frontal pole and cerebellum brain regions of AD patients (Fig. 5C). Our 2-month-old FAD cultures exhibited expression profiles of neuronal genes remarkably similar to human AD for multiple brain regions.

Similarly, we examined the expression of differentially deregulated genes in 4.5 months old FAD cultures across AMP-AD modules. There were only 5 genes that were differentially upregulated in the 4.5-month-old FAD cultures compared to controls, and these genes mainly exhibited reduced expression in some of the AMP-AD modules (Fig. S4 A). Most of the differentially downregulated genes (fold change < 1.5) in 4.5 months old FAD cultures showed reduced expression in neuronal-associated AMP-AD modules in Consensus Cluster C, except in CBEyellow and FPyellow modules (Fig. S4 B). These results suggest that neurodegeneration-related genes were downregulated in our FAD cultures, similar to human AD patients.

## Discussion

We report here that iPSC-derived NPCs reliably integrate into the silk-collagen scaffold model, expand, and differentiate in healthy neurons and astrocytic-like cells as indicated by the expression of neuronal and astrocytic markers. Accordingly, gene expression and cell enrichment analysis revealed a similar cellular composition across all cultures. More importantly, 3D neural cultures from APP London mutation heterozygous carriers recapitulate some AD-like phenotypes. We found that FAD cultures displayed an elevated Aβ42/40 ratio in conditioned media at 2 and 4.5 months, as previously reported in 2D cultures derived from the same FAD lines [15, 23]. While the Aβ42/40 ratio change was similar at the two time points, extracellular deposition of Aβ42 developed only in the older cultures and identified FAD from control cultures. Notably, Aβ42/40 ratio and Aβ42 extracellular deposition in the isogenic control cultures were similar to BR24 and YZ1 lines. These data demonstrate that the APP V7171I mutation causes such phenotypes. LFP recordings showed that our 3D cultures were electrically active starting at 2 months. While no difference in LFP frequency was observed at 2 months, by 4.5 months, FAD samples displayed significantly higher spontaneous and evoked potentials relative to controls. The increased excitability in FAD cultures correlated with extracellular Aβ42 deposition but not with soluble Aβ42/40 ratio levels, as they were similar at both time points. These data suggest that extracellular Aβ deposition may trigger enhanced network activity. Our data agree with previous work showing increased neuronal excitability in AD patients at an early stage of the disease [16, 43].

A large study conducted on carriers of FAD mutations (The Dominantly Inherited Alzheimer Network (DIAN) consortium) showed that changes in cerebrospinal fluid (CSF) biomarkers, cerebral amyloid deposition, and brain metabolism begin at least two decades before the predicted onset of clinical symptom [52]. Similarly, increased brain activity may precede AD diagnosis by 30 years in PSEN1 mutation carriers [53, 54]. More recently, neuronal hyperexcitability was described in organoids generated from isogenic iPSC lines carrying the FAD PSEN1 ΔE9, PSEN1 M146V, or APP Swedish mutations [55]. However, this is the first report of enhanced excitability in 3D cultures from patient-derived iPSC lines carrying the APP London mutation.

Differential expression analysis and GSA revealed that multiple gene sets were deregulated in FAD compared to control cultures. At 2 months, upregulated genes were associated with Cytokines, Activated Microglia, and Neuronal Cytoskeleton pathways in GSA. The downregulated genes were related to multiple neurodegeneration pathways such as Transmitter Release, Neural Connectivity, and Vesicle Trafficking. Some of the differences were maintained for up to 4.5 months. At 4.5 months, GSA identified increased expression of genes associated with Cytokines and reduced expression of genes associated with multiple pathways such as Transmitter Release, Neuronal Cytoskeleton, and Vesicle Trafficking. While most DEGs in the FAD cultures are expressed in neurons, genes associated with activated microglia were upregulated in FAD cultures at 2 months. Even though only scores for neuronal and astrocyte markers reached a statistically significant threshold in the nCounter^®^ Cell Type Profiling Module, TMEM119-positive cells are present in our cultures. Similarly, mesoderm-derived progenitors, from which microglia originate, and mature microglia-like cells innately develop in cerebral organoids [49, 50].

To validate our cultures as an AD model in vitro, we applied a system biology approach developed to evaluate AD mouse models [51]. A correlation analysis was performed between gene expression changes in FAD cultures and changes in human AD cases in post-mortem co-expression modules. We used a recent human molecular disease catalog based on harmonized co-expression data from three independent postmortem brain cohorts (ROSMAP, Mayo, Mount Sinai Brain bank) [56-58] and seven brain regions that define 30 human co-expression modules and five consensus clusters derived from the overlap of those modules [17]. FAD cultures at 2 months showed significant positive correlations with human co-expression modules in Consensus Cluster B, including transcripts enriched for immune-related pathways. They showed significant positive correlations with multiple modules in Consensus Cluster C, including transcripts enriched for neuronal-related pathways. However, these significant positive correlations diminish at 4.5 months of age.

Next, we examined the expression profile of differentially regulated genes in the FAD cultures identified by the ROSALIND^®^ platform across human AMP-AD transcriptomic data. We determined that most of the differentially upregulated genes in 2-months-old FAD cultures showed increased expression in immune/inflammation-associated AMP-AD modules in Consensus Cluster B. Similarly, most of the differentially downregulated genes in 2-months-old FAD cultures showed reduced expression across numerous human brain regions in Consensus Cluster C. Thus, 2-month-old FAD cultures exhibited expression profiles of neuronal genes remarkably similar to human AD for multiple brain regions. In the 4.5-month-old FAD cultures, most differentially downregulated genes showed reduced expression in neuronal-associated AMP-AD modules in Consensus Cluster C. These results suggest that neurodegeneration-related genes were downregulated in FAD cultures, similar to human AD patients.

Since iPSC reprogramming rejuvenates cells and erases donor aging signatures, concerns were raised about the ability to model age-related diseases with iPSC-derived neuronal cultures [59]. It is well-established that AD develops over decades [60]. Thus, genetic factors may produce endophenotypes early in life; therefore, they can be captured in iPSC-derived neural cultures.

## Conclusions

We developed a neural tissue model that supports the differentiation of iPSC-derived NPCs in a donut-shaped porous silk sponge with an optically clear (collagen-filled) central region. The porous design facilitates nutrients and oxygen diffusion to meet the metabolic demand of long-term cultures. This study provides compelling evidence that our 3D bioengineered model from patient-derived FAD iPSCs develops time-dependent phenotypes and transcriptomic features observed in AD patients. Some of these changes have been identified across other 3D models [1]. However, to our knowledge, none of these studies could recapitulate several different AD-related hallmarks in one single model. While Aβ extracellular deposition and changes in the Aβ42/40 ratio represent frequent findings in other 3D models, the enhanced excitability and transcriptomic changes described in our manuscript have been difficult to recapitulate in these conditions. Importantly, in our scaffolds, these phenotypes have been identified at two time points (2 and 4.5 months), resembling some of the features that characterize AD progression. Aβ42/40 ratio and transcriptomic changes are already present at 2 months, while Aβ deposition appears only at 4.5 months when the neuronal hyperexcitability becomes evident, suggesting a temporal relation between these two events. Gahtak et al. [55] detected hyperexcitability in AD organoids accompanied by increased Aβ immunostaining and an increase in the ratio of secreted Aβ42/40 at 2 months; nevertheless, no other time points were analyzed, providing no information on the temporal relation between these events. To date, no other studies have been published examining the transcriptomic features of these 3D models while comparing them to those of human AD brains in a large study that performed a co-expression meta-analysis of harmonized data from Accelerating Medicines Partnership for Alzheimer’s Disease (AMP-AD). These data are, in our opinion, key to understanding the pertinency of these models for studying AD-related pathomechanisms over time. As discussed, one of the main strengths of this model is that every element was designed to be tunable and amenable to experimental modification. Although we did not incorporate a layered structure for this study, the 3D model can be designed using a concentric toroidal scaffold to more closely approximate gross features of the layered brain tissue [11]. However, we need to consider that any modification to the structural design of the model could cause unexpected changes to the integrity and longevity of the cultures [14] and therefore require extensive and costly troubleshooting. We could also incorporate embedded electrodes for continuous, long-term recordings of electrophysiological activity [61, 62]. Moreover, unlike self-organizing organoids, mixed 3D neuronal tissue could be generated by co-culturing neurons, astrocytes, and microglia derived from different donors or isogenic iPSC lines carrying AD-linked genetic variants [63]. For example mixed 3D cultures containing mutant neurons and wild-type astrocytes or microglia cells and vice-versa could be used to study the impact of cell specific genetic variants on AD-like phenotypes. For all these reasons, our bioengineered neural tissue represents a unique tool to model AD-related pathomechanisms over time, with several advantages and potential compared to the existing model.

On the other hand, our model still has technical limitations. Even though the porous structure of the scaffolds facilitates the diffusion of oxygen and nutrients, it lacks a proper neurovascular unit, a perivascular compartment, and coordinated drainage of accumulated metabolites. In addition, other practical challenges like scalability and high throughput screening compatibility will have to be optimized.

## Methods

### iPSC lines

hFAD-1, hFAD-2 Clone A and B iPSC lines [15] were generated in the laboratory of Dr. T. Young-Pearse (Brigham and Women’s Hospital, Boston, MA, USA). BR24 iPSC line (ROSMAP) [9, 28] was generated at the New York Stem Cell Foundation. YZ1 iPSC line was obtained from the University of Connecticut-Wesleyan Stem Cell Core (Farmington, CT, USA [15]). The original manuscripts cited above describe the reprogramming methods and quality control assays for these lines.

hFAD-2 Clone B Corrected was generated at the University of Connecticut-Wesleyan Stem Cell Core (Farmington, CT, USA). Briefly, the guide RNA used for APP editing was designed using chop-chop (https://chopchop.cbu.uib.no/) for specificity to the loci of interest. The resulting guide 5’ GACAGTGATCATCATCACCT 3’ was identified to have high specificity to the desired loci. The guide is designed to cut the mutant allele. Given that the CAS9/guide RNA was transiently expressed in the cells, the potential for off-targets in the nearest matches was minimal. Potential off-targets were identified using the Cas-OFF finder (http://www.rgenome.net/cas-offinder/). The top eight identified sites were amplified from the clones, and Sanger sequenced to verify the status of these potential off-targets. None of the predicted sites showed evidence of off-targeting cutting. After targeting, the corrected allele resulted in a single base difference from the original guide sequence. None of the clones sequenced exhibited off-target cutting at the predicted sites or the corrected allele.

All iPSC lines have been confirmed by sequencing before starting any of the experiments listed in this manuscript.

### ROSMAP study participants

iPSC lines were generated from autopsied participants in either the Religious Orders Study or Rush Memory and Aging Project (ROSMAP) [28]. Both studies were approved by an Institutional Review Board of Rush University Medical Center; they enroll older persons without dementia who agree to annual clinical evaluation and brain donation at death. All participants signed informed consent, an Anatomical Gift Act, and a repository consent to allow their resources to be repurposed. As part of the annual evaluation, peripheral blood mononuclear cells were drawn and cryopreserved.

### Karyotype analysis and mycoplasma testing

G-banded karyotyping was outsourced to WiCell (Madison, WI, USA). WiCell’s chromosome analysis is optimized for pluripotent stem cells. All results are reviewed by an American Board of Medical Genetics and Genomics (ABMGG) board-certified or board-eligible director. Karyograms for all the lines have been checked every ten passages.

Mycoplasma testing was performed monthly for all the lines included in this study at any differentiation stage (LookOut Mycoplasma PCR Detection Kit, Sigma). All cells used in this study were consistently negative.

### NPC differentiation

iPSCs were maintained in feeder-free conditions on plates coated with Matrigel hESC-qualified Matrix (Corning) with daily media changes using mTeSR1 medium (StemCell Technologies). Cells were held at 37°C and 5% CO_2_ and were split as necessary based on colony growth (5-7 days). Cells were passaged in small aggregates (1:7/1:10 depending on the line) using ReLeSR (StemCell Technologies). 10 μM Y-27632 (ROCK inhibitor, StemCell Technologies) were added to the culturing media for a maximum of 24 hours after passage.

According to the StemCell Technologies method, NPCs were differentiated using an EB-based protocol without SMADi supported by STEMdiff Neural Induction medium (StemCell Technologies) with daily media change over ∼20 days. To generate iPSC-derived EBs for neural induction AggreWell 800 (StemCell Technologies), 24-well plates were used. These plates enable the easy generation of uniformly sized EBs, making differentiation experiments more reproducible. For each well of the AggreWell plate, 3×10^6^ single-cell suspension (Gentle Cell Dissociation Reagent, StemCell Technologies) of iPSCs were plated, followed by centrifugation at 100xg for 3 minutes, to capture the cells in the microwells. A daily partial-medium change was performed for 4 days. On day 5, EBs were harvested from a single well of an AggreWell 24-well plate and re-plated onto a Matrigel hESC-qualified-coated well of a tissue culture-treated 6-well plate. Replated EBs were maintained with daily media change until neural rosette structures were visible and defined. Around days 10-12, based on the different lines, rosettes were selected (Neural Rosette Selection Reagent, StemCell Technologies) and replated onto Matrigel hESC-qualified Matrix-coated plates. Selected rosette-containing clusters attach, and NPC outgrowths will form a monolayer between the clusters. A daily full-medium change was performed. NPCs are ready for passage 1 at approximately day 17-19 (Accutase, StemCell Technologies). NPCs were plated at 1.25×^5^ cells/cm^2^ on Matrigel hESC-qualified Matrix-coated wells and maintained in Neural Progenitor medium (StemCell Technologies) with daily complete media change. Single-cell suspensions (2×^6^ cells/mL) of NPCs were cryopreserved and banked for future usage using

Neural Progenitor Freezing (StemCell Technologies). At least two separate differentiations of NPCs have been generated from different iPSC stocks for all the lines included in this study and used to differentiate hNeurons.

### Scaffold preparation, seeding, and differentiation into hNeurons

Silk protein was processed from *Bombyx mori* cocoons as described previously [64]. In brief, silk cocoons were cut into fragments and boiled for 30 minutes in a 0.02 M Na_2_CO_3_ solution. The fibroin was solubilized in a 9.3 M LiBr solution dialyzed against deionized water. The silk solution was then diluted to 6% weight/volume and used to generate the porous scaffolds via salt leaching [19], using 500–600 um NaCl crystals to generate a sponge-like structure. A biopsy punch was used to generate the donut-shaped scaffolds (5mm outer diam. x 2mm central hole diam. x 2mm ht.). Scaffolds were autoclaved and then coated with poly-L-ornithine (20ug/mL in DPBS, Sigma) and laminin (10ug/mL in DMEM/F-12, Sigma) overnight at 4°C.

A strained (40um nylon mesh) single-cell suspension of (2×^6^/100 uL) NPCs between passages two and six was added to the coated scaffold in a 96-well plate and incubated for 24 hours. The media (200uL, Neural Progenitor medium) was changed daily for five days following seeding, moving the scaffolds to fresh wells with each change. After allowing five days of cellular expansion, the scaffolds were moved to a clean well and infused with a 100 uL solution of cold Collagen Type-1 Rat Tail (3.0mg/mL, Corning) mixed with 10X PBS and 1N NaOH (ratio 88:10:2). The collagen-filled scaffolds were then incubated at 37°C until the gelation occurred (∼45 minutes). Once solidified, the scaffolds were transferred to a 24-well plate and flooded with 1.5 mL of BrainPhys media (StemCell Technologies) supplemented with SM1 (StemCell Technologies) and N2 (ThermoFisher) with media changes every four days and differentiated into hNeurons.

### Immunofluorescence and imaging analysis

2D and 3D cultures were fixed with a 4% paraformaldehyde/4% sucrose solution in DPBS for 15 and 30 minutes, respectively, followed by membrane permeabilization with 0.3% Triton X-100 for 15 minutes. Non-specific binding sites were blocked for one (2D) or four (3D) hours at RT (5% BSA, 0.1% Triton-X-100 in DPBS). Cultures were then incubated with primary antibodies (see Antibody Table) diluted in 1% BSA, 0.1% Triton-X-100 in DPBS overnight at 4°C. After multiple washes in DPBS, sections were incubated with Alexa (1:1000 in 1% BSA in DPBS) fluorescent secondary antibodies against the corresponding host species for one (2D) or three (3D) hours at RT. Nuclei were stained with DAPI (Sigma). 2D cultures were mounted on superfrost slides (Fisher Scientific) using Fluoromount-G1 mounting media (Southern Biotech), while 3D cultures were imaged directly.

NeuroFluor™ NeuO (StemCell Technologies), a membrane-permeable fluorescent probe that selectively labels iPSC-derived neurons, was used to stain live cultures according to manufacture instructions.

Z-stack fluorescence images were obtained using confocal microscopes (SP8 Falcon Leica, SP8 Leica, see figure legends for details), keeping constant imaging parameters. 3D reconstructions were generated with the LAS X Life Science Leica software. Image analysis was performed using Fiji ImageJ version: 2.0.0-rc-69/1.52p.

Aβ burden volumetric analysis was performed using the Fiji ImageJ plug-in 3D object counter. The sum of all extracellular Aβ deposition volumes was divided by the total volume of the region of interest (ROI) imaged to obtain the amyloid burden values. At least two separate sets of cultures per line and three images per set of cultures have been analyzed.

### MSD assay and data analysis

Before harvest, BrainPhys media was changed, and fresh media was dispensed into each well. Twenty-four hours later, media was collected and stored at -80°C. MSD (Rockville, MD, USA) assay for the detection of human Aβ38, Aβ40, and Aβ42 (V-PLEX Aβ Peptide Panel 1 (4G8) Kit) was performed in-house according to the manufacturer’s instructions on undiluted conditioned media. Technical duplicates were run for all the samples analyzed. The assay demonstrated high sensitivity with a lower limit of detection values in the low-pg/mL and over 2.5 logs of dynamic range, allowing the measurement of analytes in all samples, except for Aβ38, which fell under the lower limit of detection for most of the samples. All MSD assays exhibited excellent reproducibility with an average calculated concentration of 3.6% and 7.2% for the standards and samples within the detection range of the assay, respectively. The data were analyzed using MSD Workbench 4.0 software. The software fits the standard curves using a four-parameter logistic fit with 1/y^2^ weighting.

### LFP recordings

Local field potentials (LFPs) were recorded from samples, adapting a protocol by Du et al. [37]. Both spontaneous and evoked potentials (EPs) were collected. Each sample was placed in the center of a 35mm plastic petri dish (Corning) containing 2 mL of an extracellular solution simulating cerebrospinal fluid (mM): 130 NaCl, 1.25 NaH_2_PO_4_, 1.8 MgSO_4_, 1.6 CaCl_2_, 3 KCl, 10 HEPES-NaOH, 5.5 glucose, pH 7.4. The dish and solution were maintained at 37°C using a WP-16 Warmed Platform (with a TC-134A Handheld Temperature Controller). A bipolar stimulator was positioned on top of the sample, affixing it to the bottom of the dish. Then, a recording electrode (borosilicate glass pipette, 40-80 MΩ resistance) which was pulled with a Sutter P-97 (Navato), was inserted into the silk-scaffold region between the stimulators (e.g., stimulator-electrode-stimulator). LFPs were amplified and digitized (Axon Instruments DAC; Intan digital amplifier) with a sampling frequency of 2500Hz. Recordings were performed blind (each sample was given an unidentifiable ID), always counterbalanced by condition, and replicated batches were measured within the same 12-hour window.

Spontaneous activity (baseline) was recorded for 2 mins, followed by the evoked potential phase, which consisted of a 60s period of stimulation (1ms pulse, 1Hz, 5 mA), an injection of 10 µM tetrodotoxin (sodium channel blocker), 2 mins of perfusion, and a final 60 sec period of stimulation. Data were retrieved from Clampex 10.7 (Axon Instruments), exported to Clampfit 10.7, and “spikes” indicative of activity were extracted using an amplitude threshold (±0.35 mV). Spike frequencies (LFP spikes/minute) were computed for spontaneous (baseline) recordings. Evoked potentials per stimulation period were also quantified after a background-subtraction process as previously described [37]. All data were normalized to DNA content quantified using Quant-iT PicoGreen dsDNA Assay kit (ThermoFisher). Data were also subjected to a log-10 transformation to stabilize and normalize expected trial-to-trial variation.

### RNA isolation and NanoString outsourcing

At the time of desired analysis, scaffolds were snap-frozen using liquid nitrogen and stored at −80°C. The frozen scaffolds were then homogenized using a liquid nitrogen-chilled Spectrum Bessman Tissue Pulverizer (Fisher Scientific). RNA from homogenized samples was extracted immediately after the homogenization using the RNeasy Plus Mini kit (Qiagen) following the manufacturer’s instructions. Briefly, samples were dissolved in Qiagen RLT Plus lysis buffer supplemented with 1% β-mercaptoethanol and centrifuged for 3 min at maximum speed. The supernatant was transferred to a gDNA Eliminator spin column to remove genomic DNA, and the flow-through was mixed with 70% ethanol and placed in an RNeasy spin column. After one wash with the RW1 buffer and two washes with the RPE buffer, RNA was eluted in 20uL of RNase-free water and quantified using a NanoDrop 1000 Spectrophotometer (ThermoFisher). RNA concentration was normalized to 20ng/uL, and 150ng of total RNA was shipped to NanoString Technologies^®^ (Seattle, WA, USA) for analysis.

### NanoString neuropathology panel gene expression profiling

The NanoString Neuropathology profile panel was used for gene expression profiling of our RNA samples on the nCounter platform (NanoString Technologies^®^, Seattle, WA, USA). The neuropathology panel incorporated 770 human genes, including 10 references (housekeeping genes) and 760 endogenous genes, to target all aspects of neurodegeneration. Data were analyzed by ROSALIND® (https://rosalind.onramp.bio), with a Hyperscale architecture developed by ROSALIND, Inc. (San Diego, CA, USA). Normalization, fold changes, and p-values were calculated using criteria provided by NanoString. ROSALIND® follows the nCounter® Advanced Analysis protocol of dividing counts within a lane by the geometric mean of the normalizer probes from the same lane.

Housekeeping probes for normalization were selected based on the geNorm algorithm implemented in the NormqPCR R library [65]. Fold changes, and significance score (p-value) were calculated using the optimal method described in the nCounter® Advanced Analysis 2.0 User Manual. Significant p-values (p < 0.05) were adjusted for multiple genetic comparisons using the Benjamini–Hochberg method of estimating false discovery rates [66]. Gene expression was considered significantly different if there was a greater than 1.5. fold change compared to controls. The nCounter Neuropathology profile panel annotation database was referenced to perform gene set enrichment analysis. Directed global significance scores (GSS) were calculated relative to a set of background genes measured in the experiment. This analysis determined both significance and directional response (i.e., upregulated or downregulated) of GSS for a given neurodegeneration pathway. A total of 24 fundamental neurodegeneration pathways were examined to determine the effect of FAD mutation. Bar plots were generated to compare neurodegeneration pathways as a function of directed GSS.

### AMP-AD post-mortem brain cohorts and gene co-expression modules

Data on the 30 human AMP-AD co-expression modules were obtained from the Synapse data repository [56, 58, 67] (https://www.synapse.org/#!Synapse:syn11932957/tables/;SynapseID: syn11932957). These modules were obtained from a co-expression meta-analysis of harmonized AMP-AD data across three independent study cohorts. Each module is enriched in genes differentially expressed in AD cases versus controls [17]. The 30 AMP-AD modules were further grouped into five consensus clusters that describe the major functional groups of alterations observed in human AD [17, 51].

### iPSC-derived cultures-human brains expression comparison

The nSolver software-generated raw gene expression counts for the Neuropathology panel. Normalization was done by dividing counts within a lane by the geometric mean of the housekeeping genes from the same lane. Next, normalized count values were log-transformed for downstream analysis. To determine the effect of FAD mutation, we fitted a linear mixed-effect regression model using the lme4 function in R by considering the fixed effect of FAD mutation plus random variation in intercept among cell lines within controls and FAD mutation (formula ∼ FAD + 1|FAD:Cell.Line).

To compare expression changes in FAD lines with those observed in human disease, we computed Pearson correlations between log fold change in transcript expression of human AD patients compared to control patients (Log2FC(AD/controls) and the effect of FAD mutation estimated by the linear mixed-effect regression model [51, 68]. Correlation coefficients were computed using the cor.test function built-in R as: cor.test(log2FC(AD/Control,β_FAD_).

LogFC values for human transcripts after correction for technical covariates were obtained via the AD Knowledge Portal (https://www.synapse.org/#!Synapse:syn14237651).

### Statistical analysis

Information regarding statistical analyses can be found in the figure legends. Data are expressed as the mean ± standard error of the mean (SEM), represented as error bars. All statistical tests were computed using GraphPad Prism 9 software unless otherwise noted. ROUT method (1%) was applied to identify outliers. A p-value of 0.05 was used as the significance threshold throughout this study. In all figures, p-values are illustrated for all tests used: ^n.s.^ p > 0.05, * p < 0.05, ** p ≤ 0.01, *** p ≤ 0.001, **** p ≤ 0.0001. A minimum of two differentiations for each iPSC line have been analyzed for all experiments. Figure legends and graphs representing individual data points indicate the sample size (n = number of scaffolds).

## List of supplementary materials

Fig. S1, Fig. S2, Fig. S3, Fig. S4, Table 1.

**Table 1:** List of antibodies used throughout the study.

| Target Antigen | Host | Clone | Clonality | Reactive Species | Company | Catalog # | IF Dilution |  |
| --- | --- | --- | --- | --- | --- | --- | --- | --- |
|  |  |  |  |  |  |  | 2D | 3D |
| **Aβ42** | Rabbit | H31L21 | Monoclonal | Human, Mouse | Invitrogen | 700254 | / | 1:200 |
| **GFAP** | Rabbit | / | Polyclonal | Human<br>90-95% homology<br>between species | Dako | Z0334 | 1:1000 | 1:500 |
| **GFAP** | Mouse | G-A-5 | Monoclonal | Pig, Rat, Human | Sigma | G3893 | / | 1:500 |
| **MAP2B** | Mouse | 18 | Monoclonal | Rat, Human,<br>Mouse | BD Biosciences | 610460 | 1:1000 | 1:500 |
| **MAP2B** | Chicken | / | Polyclonal | Human, Rat,<br>Bovine | Millipore | AB5543 | 1:1000 | 1:500 |
| **Nestin** | Mouse | 10C2 | Monoclonal | Human<br>Non-Human Primate | StemCell Tech | 10C2 | 1:1000 | / |
| **NFH** | Chicken | / | Polyclonal | Human, Rat, Mouse,<br>Bovine, Porcine, Feline | Millipore | AB5539 | / | 1:500 |
| **OCT4** | Mouse | 3A2A20 | Monoclonal | Human | StemCell Tech | 60093 | 1:250 | / |
| **Synaptophysin 1** | Mouse | 7.2 | Monoclonal | Human, Rat,<br>Mouse, Mammals | Synaptic Systems | 101011 | 1:500 | / |
| **TMEM119** | Rabbit | / | Polyclonal | Human | Sigma | HPA051870 | / | 1:200 |
| **Tuj1** | Mouse | TU-20 | Monoclonal | Bovine, Rat, Avian,<br>Monkey, Sheep,<br>Human, Mouse, Pig | Chemicon | MAB1637 | 1:1000 | 1:500 |

## Availability of data and materials

All data generated or analyzed during this study are included in this published article and its supplementary information files. Additional information is available from the corresponding author on request. ROSMAP resources can be requested at https://www.radc.rush.edu.

## Authors’ contributions

The authors confirm contributions to the manuscript as follows: conceptualization: SL, GT; data curation: SL, RSP, NR; formal analysis: SL, RSP, NR; investigation: SL, NR, BM, WK, WLC; methodology: SL, RSP, NR, GWC; resources: WLC, GWC, DAB, TLYP, DLK; project administration: GT; supervision: SL, GWC, DLK, GT; validation: SL, RSP, NR, BM, WK, GT; visualization: SL, RSP, NR; writing - original draft preparation: SL, RSP, NR, GT; writing - review and editing: SL, RSP, NR, BM, WK, WLC, PGH, DAB, TLYP, GWC, DLK, GT.

## Acknowledgments

We thank Rachel Willen, Edward K. Robinson, Griffin Sigal, and Isabel Paine for their technical support.

## Abbreviations

2D: Two-Dimensional
3D: Three-Dimensional
Aβ: Amyloid Beta
AD: Alzheimer’s Disease
AMP-AD: Accelerating Medicines Partnership for Alzheimer’s Disease
AP: Action Potential
APP: Amyloid Precursor Protein
CBE: Cerebellum
CSF: Cerebrospinal Fluid
DEG: Differentially Expressed Gene
DIAN: Dominantly Inherited Alzheimer’s Network
EB: Embryoid Body
EP: Evoked Potential
FAD: Familial Alzheimer’s Disease
GFAP: Glial Fibrillary Acidic Protein
GSA: Gene Set Analysis
hNeuron: Human Neuron
IFG: Inferior Frontal Gyrus
iPSC: Induced Pluripotent Stem Cell
LOAD: Late-Onset Alzheimer’s Disease
LFP: Local Field Potential
MAP2B: Microtubule-Associated Protein 2B
MSD: Meso Scale Discovery
NFH: Neurofilament Heavy Chain
NFT: Neurofibrillary Tangles
NPC: Neural Precursor Cell
Oct4: Octamer-Binding Transcription Factor 4
PHG: Parahippocampus Gyrus
PSEN1: Presenilin 1
PSEN2: Presenilin 2
ROSMAP: Religious Order Study Rush Memory and Aging Project
SYN: Synaptophysin
TCX: Temporal Cortex
Tuj1: Neuron-Specific Class III Beta-Tubulin

---

**Figure S1:**
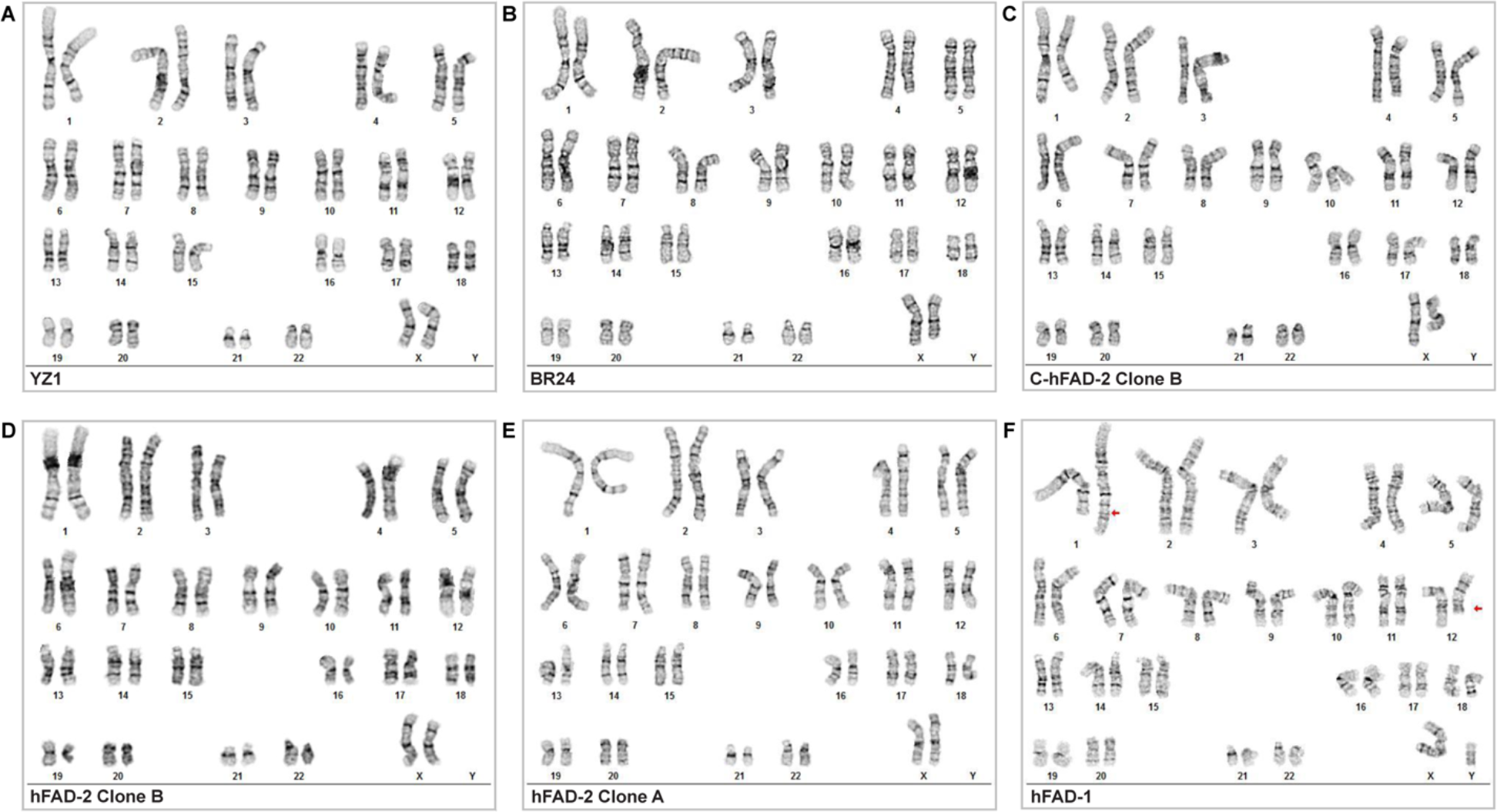
Karyotype reports for all the lines used across the study (WiCell).

**Figure S2:**
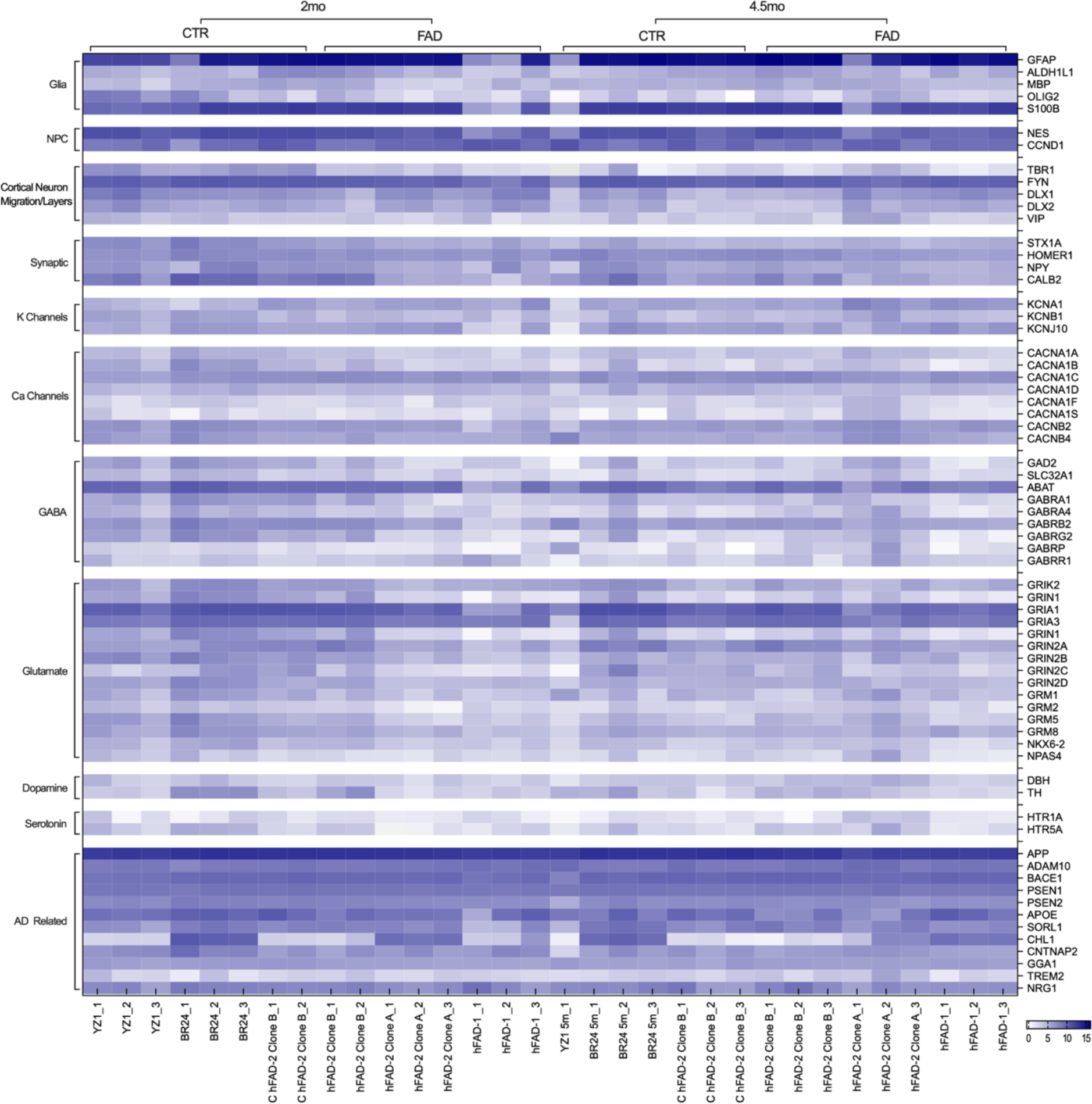
Normalized expression heatmap (9 housekeeping probes) of genes used to characterize the 3D cultures. FAD 2mo n=8, CTR 2mo n=8, FAD 4.5mo n=9, CTR 4.5mo n=7. Genes included in the heatmap are not deregulated when comparing FAD vs. CTR lines at any time point analyzed.

**Figure S3:**
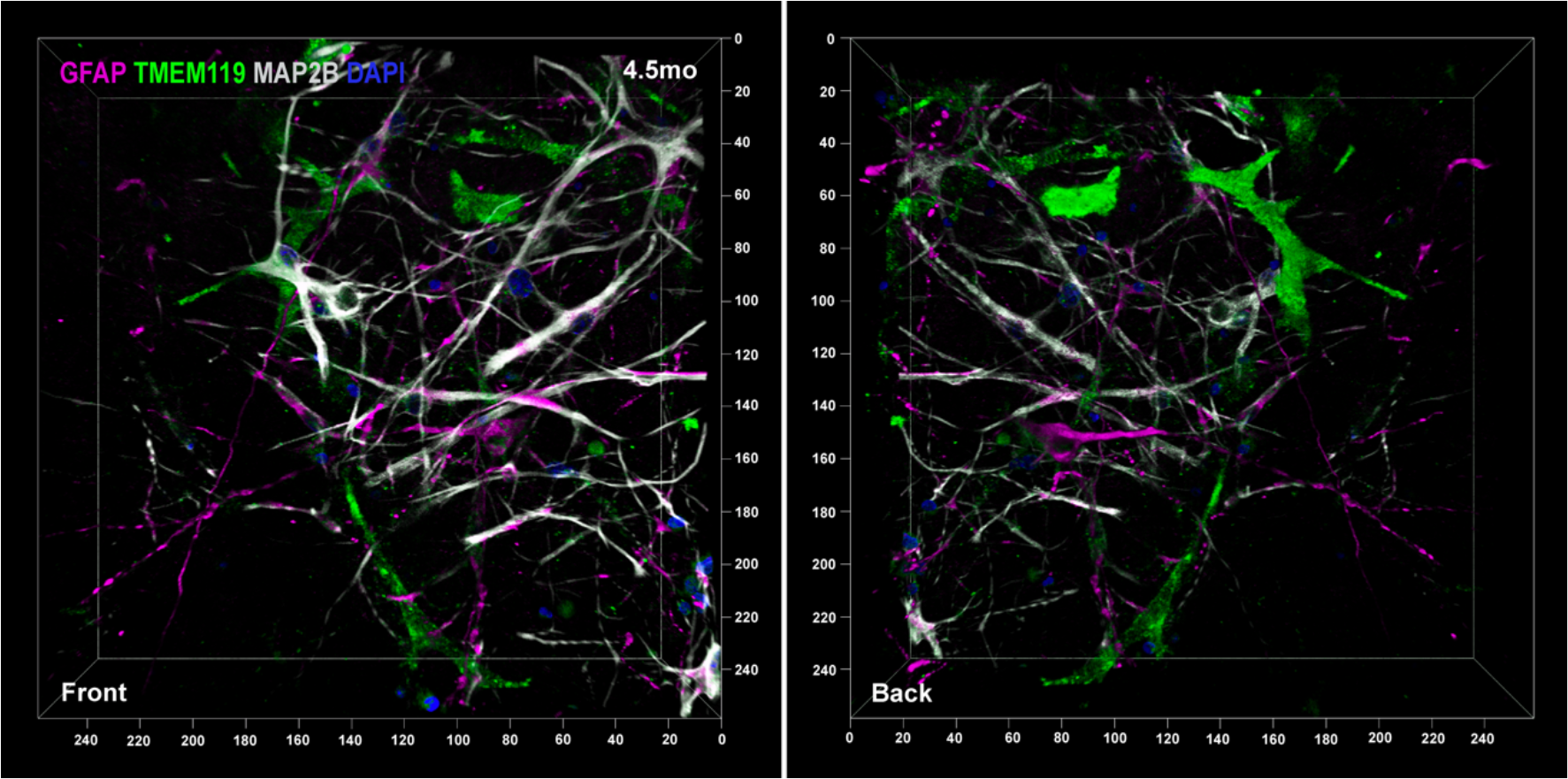
3D reconstruction of 4.5 months cultures (stained for MAP2B (grey), TMEM119 (green), GFAP (magenta), DAPI (blue)). Front and back projections are shown. Leica Falcon SP8 HC PL APO CS2 20x/0.75 IMM. 2.25x Zoom.

**Figure S4:**
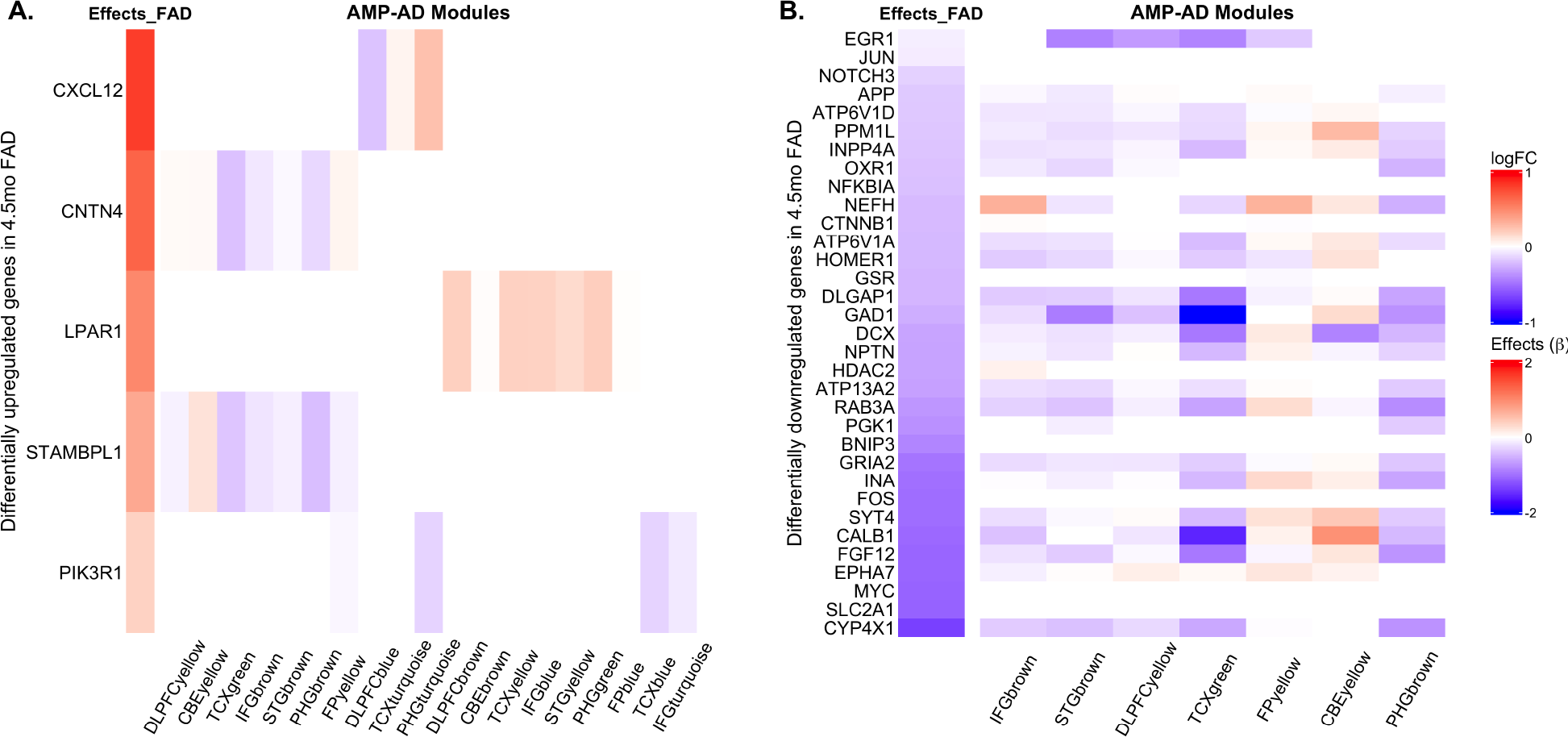
Gene expression changes of differentially expressed genes in the 4.5-months-old FAD cultures across human co-expression modules. (A) Gene expression changes of five differentially upregulated genes in the 4.5-months-old FAD lines (FDR <0.05, fold change > 1.5) across AMP-AD modules are shown in the heatmap. (B) Gene expression changes of differentially downregulated genes in the 4.5-months-old FAD lines (FDR <0.05, fold change < 1.5) across AMP-AD. FAD 4.5mo n=9, CTR 4.5mo n=7.

